# Ionic control of porin permeability in bacteria

**DOI:** 10.1101/2022.07.13.499887

**Authors:** Santiago E. Caño Muñiz, Anja Hagting, Ieuan E. Evans, Ali F Alsulami, David Summers, Tom L Blundell, R. Andres Floto

## Abstract

Bacterial porins permit permeation of hydrophilic nutrients and antibiotics across the outer membrane but also contribute to proton leak from the periplasmic space, suggesting that their activity might be dynamically regulated. Here we show, in *Escherichia coli*, that porin permeability is controlled by changes in periplasmic ions, inhibited by periplasmic acidification, thereby limiting proton loss during electron transport chain activity, and enhanced during starvation, promoting nutrient uptake. Growth in glucose increases periplasmic potassium through activating the voltage-gated channel Kch, triggering enhanced porin permeation and membrane action potentials. This metabolic control of porin permeability explains the recognized decrease in antibiotic susceptibility when bacteria are grown in lipid media and the impact of mutations in central metabolism genes on drug resistance, identifying Kch as a therapeutic target to improve bacterial killing by antibiotics.

**One sentence summary:** The permeability of bacterial porin is dynamically regulated by periplasmic pH and potassium levels, altering antibiotic resistance.

## Main text

The outer membrane of gram-negative bacteria forms a physical and mechanical barrier that protects them from chemical and biological attacks (*1–3*). Bacterial porins are water-filled β barrel channels across this membrane (*4*), mediating permeability to nutrients, such as glucose (*5, 6*), and to many antibiotics (*7*), including β lactams (*8–10*), carbapenems (*11, 12*), and fluoroquinolones (*13, 14*).

However, porin channels would also be expected to cause proton leakage from the periplasm (*15*), thereby dissipating the proton motive force generated by the electron transport chain that is required for ATP synthesis via oxidative phosphorylation (*16–20*) and other cellular processes (*18*), such as active solute transport, drug efflux (*21*), and flagellar motion (*22–24*). We, therefore, wondered whether bacteria might dynamically regulate porin channel activity, thereby balancing nutrient uptake, antibiotic susceptibility, and energy production.

We first employed the fluorescent glucose analogue 2-deoxy-2-[(7-nitro-2,1,3-benzoxadiazol-4-yl)amino]-D-glucose (2NBDG), whose entry into *E. coli* is concentration and time-dependent (***Figure 1A, B***) and known to be mediated by porins (*25–29*), to screen for bacterial knockout mutants (from the KEIO collection (*30*)) that have reduced porin activity. We observed decreased 2NBDG accumulation in mutants with deletions in porins (*ompF, ompC, ompG, nmpC*, and *phoE*) as expected, but surprisingly also in mutants for several ion channels, including the putative voltage-gated potassium channel *kch* (*31*) (***Figure 1C***); findings that were mirrored using other tracers of porin permeability (such as the fluorescent penicillin analogue Bocillin FL (*32, 33*) and Hoechst (*34*); ***Figure S1***).

**Figure 1.**
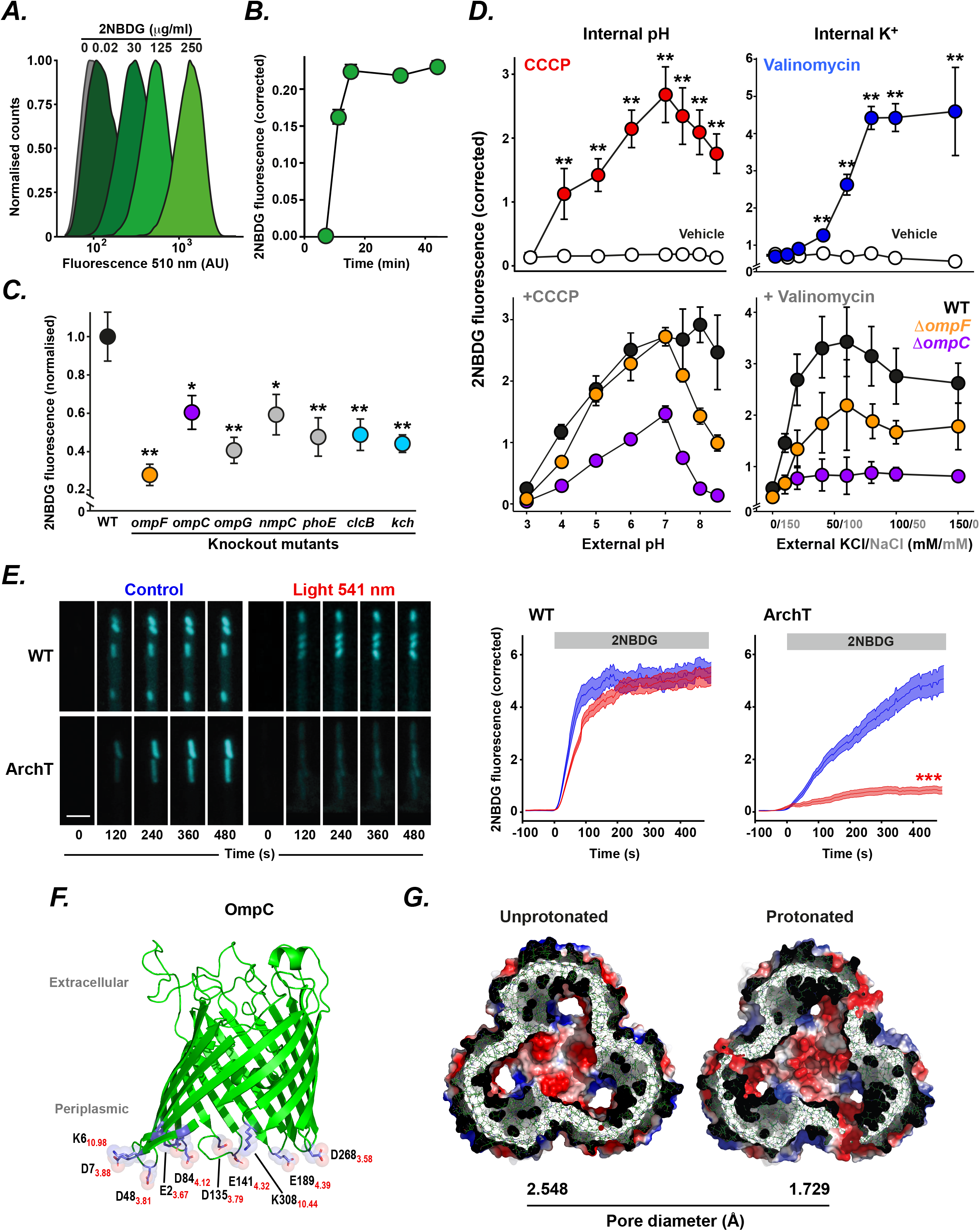
Bacterial porins are regulated by changes in internal proton and potassium concentrations. (**A, B**) Accumulation of the fluorescent glucose analogue 2-Deoxy-2-[(7-nitro-2,1,3-benzoxadiazol-4-yl)amino]-D-glucose (2NBDG) into *E. coli* (**A**) following 10 minute incubation at a range of concentrations and (**B**) at 20 mg/ml over a range of incubation times. (**C**) 2NBDG accumulation following incubation for 10 min at 20 µg/mL in wild-type *E. coli* (black) or isogenic knockout strains for the major porins *ompF* (*yellow*) and *ompC* (*purple*), minor porins (*ompG, nmpC, phoE*; *grey*), the chloride channel *clcB* (*blue*) and the voltage-gated potassium channel *kch* (*blue*). (**D**) Effect on 2NBDG accumulation in *E. coli* of (*left upper panel*) changing external pH alone (*white*) or in the presence of CCCP (250 μM) concentration; *red*); (*right upper panel*) changing external potassium concentrations (while maintaining monovalent cations constant) alone (*white*) or in the presence of valinomycin (100 mM; *blue*); (*lower panels*) changing external pH in the presence of CCCP (*left*) or changing external K+ in the presence of valinomycin (*right*) on wild type bacteria (*black*) and ompF (*yellow*) or ompC (*purple*) isogenic knockout strains. (**A-D**) Data (mean ± SEM) are representative of at least three independent experiments performed in at least triplicate. \**p* < 0.05, ** *p* < 0.01 (Student’s t-test). (**E**) Single-cell fluorescence imaging of *E. coli* grown in a microfluidic perfusion system (*Mother machine*, (*37*)) shows 2NBDG accumulation over time in wild-type bacteria expressing empty vector (WT) or expressing the light-activated proton pump ArchT in the inner membrane in the presence (*blue*) or absence (*red*) of 541 nm light exposure. Data (mean ± SEM) are representative of at least three independent experiments performed in at least triplicate, imaging at least 50 individual bacteria per condition on each occasion. *** *p* < 0.001 (Student’s t-test). (**F**) Structure of OmpC (PDB 2J1N) highlighting periplasmic residues (E2, E43, E189, K6, K308, D7, D48, D135, D141, D268) likely to be affected by periplasmic acidification (pKa values in red). (**G**) Cross-sectional views of molecular dynamic simulations of the effect on pore diameter of protonation of residues are illustrated in (F), thereby modelling periplasmic acidification.

To examine whether porin permeability might be under ionic regulation, we monitored 2NBDG uptake in wild-type *E. coli* (using flow cytometry) across a range of external pH and potassium concentrations (while maintaining total external monovalent cations constant). We found that 2NBDG uptake was significantly enhanced as external pH increased but only in the presence of the protonophore CCCP; and as external K^+^ concentrations increased in the presence of the potassium ionophore valinomycin with effects mainly mediated by the OmpC and OmpF porins (***Figure 1D***). These findings suggest that internal (rather than external) proton and potassium levels control porin permeability.

We, therefore, explored whether directly altering periplasmic ions could influence porin permeability. We expressed the light-activated proton pump, ArchT (*35, 36*), in the inner membrane of wild-type *E. coli* to selectively acidify the periplasm and quantified 2NBDG uptake using a microfluidic perfusion system that allows imaging of individual bacteria over time (*37, 38*). We found that, upon light exposure, ArchT-expressing *E. coli* (but not controls) showed reduced 2NBDG uptake (***Figure 1E; Supplementary Movie 1***), indicating that porin permeability is reduced upon periplasmic acidification.

Periplasmic pH and K^+^ may have direct effects on the pore diameter of porins, as indicated by previous lipid bilayer electrophysiological studies that have reported pH-dependent and K^+^-dependent effects on porin conductance (*7, 39–44*); the periplasmic surface of OmpC is decorated with charged residues (***Figure 1F***) that could be affected by changes in periplasmic pH and K^+^ (via altered protonation and electrostatic shielding respectively (*43*)), and molecular dynamic simulations suggest that periplasmic acidification will reduce the pore diameter of OmpC (***Figure 1G***).

We next examined how ion concentrations within bacteria might change over time by creating genetically-encoded fluorescent sensors for cytoplasmic and periplasmic pH (based on pHlourin and pHuji, respectively (*45, 46*)) which decrease in fluorescence with acidification, and cytoplasmic and periplasmic K^+^ (based on GINKO1 and GINKO2 respectively (*47*)), and cytoplasmic Ca^2+^ (based on CaM (*48*)) which increase in fluorescence with increasing ion concentration. Periplasmic localization of sensors was achieved by exploiting the *pelB* export system (as previously described (*49, 50*)). Using single-cell imaging of bacteria, we observed dynamic changes in cytoplasmic and periplasmic H^+^ and K^+^ ions (but not cytoplasmic Ca^2+^) over time (***Figure 2A, B***), with considerable volatility seen in periplasmic H^+^ and K^+^ levels (***Figure 2C***), raising the possibility of temporal regulation of porin activity through these ion fluctuations.

**Figure 2.**
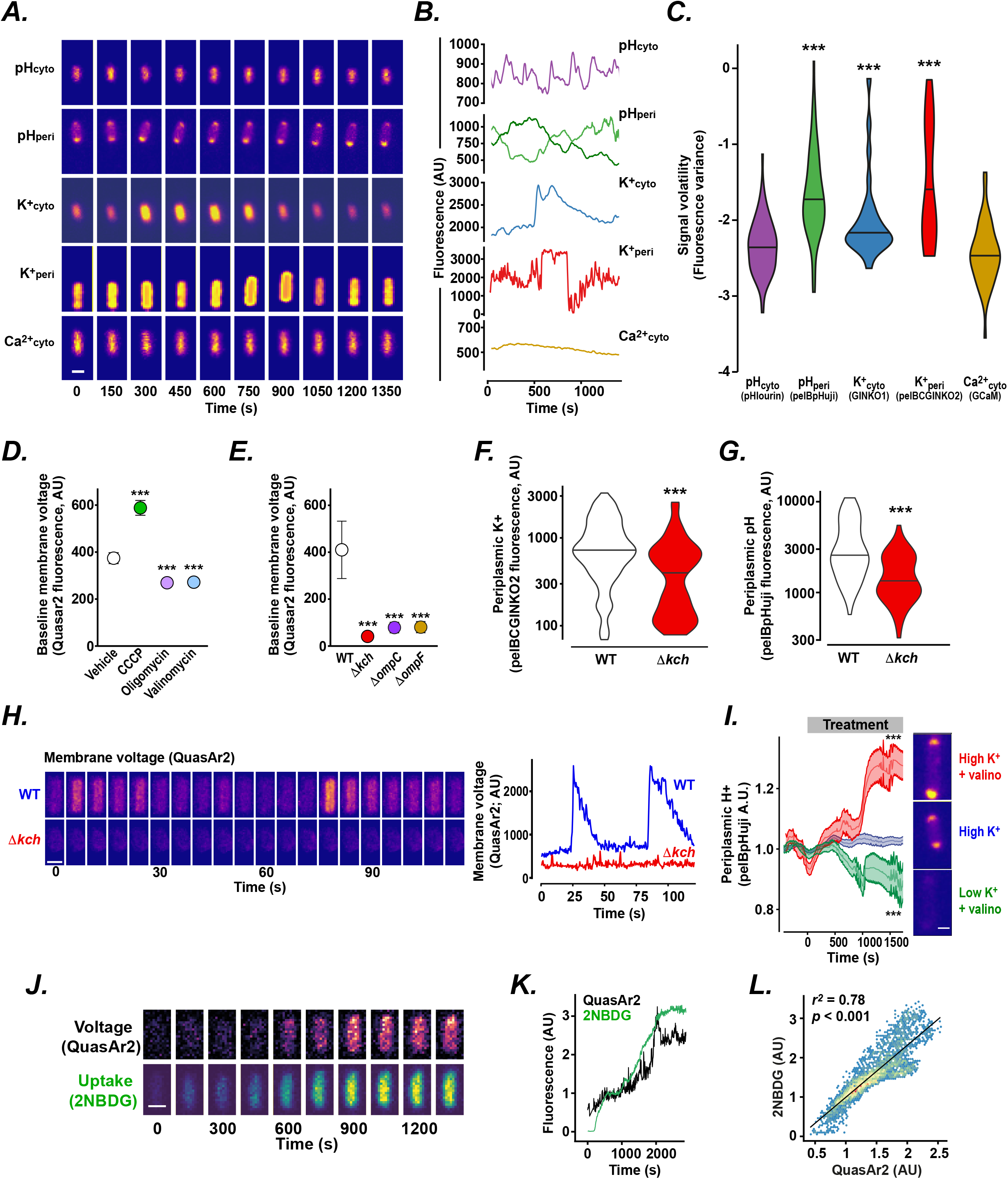
Fluctuations in periplasmic ions cause temporal changes in porin permeability. (**A-C**). Representative images (**A**) and traces (**B**) over time of different individual bacteria (growing in M9 + 1 g/L casamino acids + 1 g/L glucose and 1 mM tryptophan media) imaged in the *Mother machine* microfluidics platform expressing fluorescent sensors for cytoplasmic pH (pHcyto; pHluorin; *purple*), periplasmic pH (pHperi; pelB-pHuji: *green*), cytoplasmic K^+^ (K^+^cyto; Ginko1; *blue*), periplasmic K^+^ (K^+^peri; pelBC-ginko2; *red*), or cytoplasmic calcium (Ca^2+^cyto; GCaMp6f; *yellow*). (**C**) Signal variance over time for each sensor fluorescence value. Data are representative of at least three independent experiments performed in at least triplicate, imaging at least 30 individual bacteria per condition. \*\*\**p* < 0.001 (Student’s t-test relative to Ca^2+^cyto signal). (**D, E**) Baseline inner membrane voltage of individual bacteria (measured using the sensor QuasAr2 (*51*), which increases in fluorescence with membrane depolarisation). (**D**) Effect on baseline membrane voltage of treatment with CCCP (concentration; *green*), oligomycin (concentration; *violet*), valinomycin (concentration; *light blue*), or vehicle alone (*white*). (**E**) Baseline membrane voltage in wild-type cells (WT, *white*) or isogenic knockouts for *kch* (*red*), ompC (*purple*), or ompF (*yellow*). (**F, G**) Single-cell measurements of (**F**) periplasmic K+ and (**G**) periplasmic pH (monitored by pelBC-GINKO2 and pelB-pHuji fluorescence, respectively) in wild-type *E. coli* (*white*) or isogenic *kch* knockout strains (*red*).. (**H**) Time-lapse montage and trace of *E. coli* WT and *Δkch* cells expressing the membrane potential reporter QuasAr2. Representative images and traces of membrane voltage (measured by QuasAr2 fluorescence) in wild-type *E. coli* (*blue*) or isogenic *kch* knockouts (*red*). (**I**) Single-cell recordings of periplasmic pH over time (monitored by pelB-pHuji fluorescence) following treatment with 150 mM external K^+^ (0 mM external Na+) and 100 mM valinomycin (*red*), 150 mM external K^+^ (0 mM Na^+^) alone (*blue*), or 0 mM external K^+^ (150 mM external Na+) and 100 mM valinomycin (*green*). Representative images after 1500 s treatment shown. (D-H) Data are representative of at least three independent experiments performed in at least triplicate, imaging at least 50 individual bacteria per condition shown as (D,E,H) mean ± SEM (Student’s t-test) or (F-G) violin plots (Wilcoxon signed-rank test). \*\*\**p* < 0.001. (**J-L**) Representative images (**J**) and traces (**K**) of simultaneous recordings in single bacteria (wild-type *E. coli*) of membrane voltage (QuasAr2 fluorescence; *black*) and 2NBDG accumulation (*green*) over time. (**L**) Plot of membrane voltage and 2NBDG uptake in individual bacteria (n > 40) over time. (I-L) The data shown are representative of at least three independent experiments per condition.

Since changes in electrochemical gradients of H^+^ and K^+^ across a membrane would be expected to affect its voltage, we employed a genetically-encoded sensor (QuasAr2, (*51*)) to measure the bacterial inner membrane potential. We found that inner membrane voltage was influenced by changes in both periplasmic H^+^ and K^+^ (***Figure 2D***) since reducing periplasmic H^+^ (by treatment with CCCP) caused membrane depolarisation; increasing periplasmic H^+^ (by inhibiting ATP synthase activity using oligomycin) caused hyperpolarisation, and reducing periplasmic K^+^ (with valinomycin) caused hyperpolarisation. Furthermore, we found that membrane hyperpolarisation was also present in deletion mutants of *ompC* or *ompF* (supporting the concept of proton loss through porins (*52*) and in *Kch* knockouts (***Figure 2E***), which we found have reduced periplasmic K^+^ levels (***Figure 2F***) and increased periplasmic H^+^ levels (***Figure 2G***), consistent with decreased cytosol-to-periplasmic K^+^ flux (*53*) and a subsequent reduction in porin-mediated proton loss respectively.

On single imaging cells, we detected periodic membrane depolarisations (‘action potentials’), in wild-type bacteria, as previously described (*54*), but not in *kch* knockout cells (***Figure 2H; Supplementary Movie 2***). Since artificially altering periplasmic K^+^ levels in wild-type bacteria (by exposure to high or low potassium in the presence of valinomycin) leads to rapid changes in periplasmic H^+^ (***Figure 2I***), the initiation of action potentials may be due to Kch-mediated K^+^ influx into the periplasm leading to porin opening and subsequent proton loss. This model predicts that depolarised bacteria will have increased porin permeability and is supported by our observation of a tight correlation between membrane voltage and 2NBDG uptake (***Figure 2J-L***).

We next explored the impact of metabolism on periplasmic ions and, consequently, porin permeability. Exposure of wild-type *E. coli* to glucose led to rapid and large fluctuations in periplasmic K^+^ and H^+^ levels, not seen in minimal media or media with lipids as the primary carbon source (***Figure 3A & B***), suggesting that Kch opening might be related to the activity of the electron transport chain (ETC) and thus metabolic state of bacteria, as previously proposed (*53*). Consistent with this idea, we found that the frequency of action potentials increased with the quality and quantity of available carbon sources, pyruvate and glucose (preferred substrates for the Kreb’s cycle (*37, 55*)), triggering the greatest number of action potentials at any given concentration (***Figure 3C & D***).

**Figure 3.**
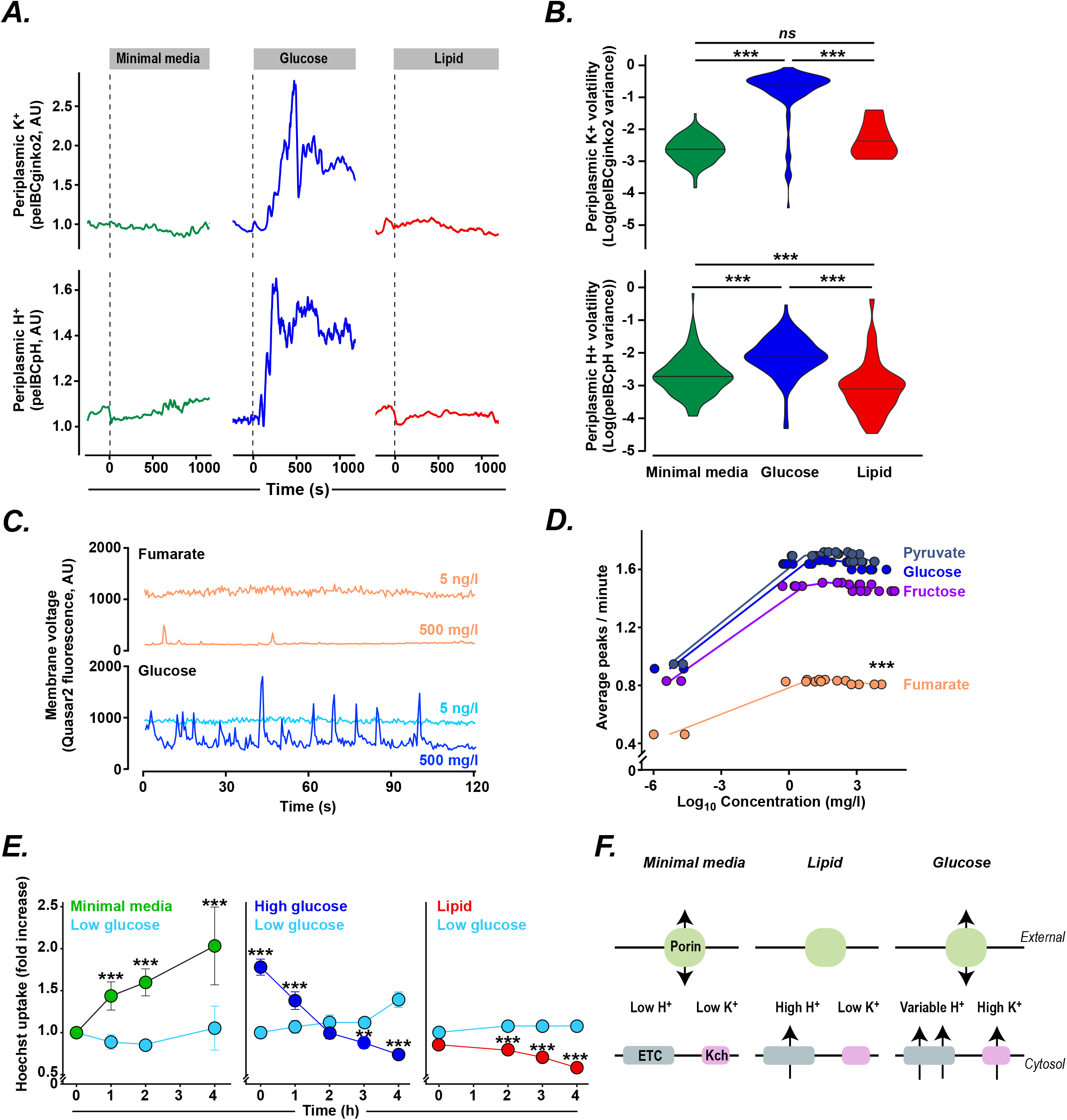
The impact of metabolism on periplasmic ions, membrane voltage and porin permeability. (**A**) Representative traces and (**B**) signal volatility (variance) of (*top*) periplasmic K^+^ and (*bottom*) periplasmic H^+^ (monitored by pelBC-GINKO2 and pelB-pHuji fluorescence, respectively) in individual bacteria when external media is changed to minimal media (M9, *green*), glucose media (M9 supplemented with 4 g/L glucose; *blue*), or lipid media (M9 supplemented with 0.014 g/L DPPC). Data shown are representative of at least three independent experiments performed in at least triplicate per condition. \*\*\**p* < 0.001 (Wilcoxon signed-rank test). (**C**) Representative traces of changes in membrane voltage over time (monitored by QuasAr2 fluorescence) in individual bacteria exposed to low (5 ng/L) or high (500 mg/L) fumarate (*pink*) or glucose (*blue*). (**D**) Frequency of action potentials (average peaks/ minute) for wild-type *E. coli* following exposure to media with different concentrations of fumarate (*pink*), fructose (*purple*), glucose (*blue*), or pyruvate (*dark blue*) as the only carbon source. Statistical analysis was performed using a generalized linear model (Supplementary methods). (**E**) Porin permeability (detected through Hoechst accumulation measured by flow cytometry) of wild-type *E. coli* exposed to M9 media alone (minimal media; *green*), M9 media with low glucose (0.04 g/L; *light blue*), high glucose (4 g/L; *dark blue*), or lipid (0.014 g/L DPPC; *red*). Bacteria were incubated for 10 minutes at 37 °C with Hoechst at times indicated. Fluorescence normalized to low glucose fluorescence at 0h. Data shown (mean ± SEM) is representative of at least three independent experiments per condition, each performed in at least triplicate. \*\*\**p* < 0.001 (Student’s t-test). (**F**) Model explaining metabolic control of porin permeability. During growth in minimal media, the permeability of porins (*green*) is high due to low periplasmic H^+^ and K^+^ levels (resulting from minimal electron transport chain (ETC; *grey*) activity and no opening of Kch channels (*pink*), respectively). Growth in lipid causes an increase in periplasmic H^+^ (through increased ETC activity) without activating Kch, resulting in low porin activity. In contrast, growth in glucose media drives ETC activity and Kch channel opening, leading to fluctuating periplasmic H^+^ and high periplasmic K^+^, causing porin permeability to increase.

As expected from these results, the metabolic state of bacteria also affected porin permeability (***Figure 3E***). We found a greater porin-mediated entry in bacteria exposed to minimal media than low glucose media (in keeping with greater porin opening under conditions of low periplasmic H^+^). We also observed greater permeability in high glucose that in low glucose media and in low glucose than in lipid media (consistent with greater porin opening under conditions of higher periplasmic K^+^).

We, therefore, propose a model for metabolic control of porin regulation (***Figure 3F***) whereby: in minimal media, porin permeability is high (due to low ETC activity and thus low periplasmic H^+^); during growth in lipid or other slowly utilized carbon sources, porin permeability is low (due to high periplasmic H^+^ and low K^+^ levels); while during growth in rich media (such as glucose), porin permeability is high (due to high periplasmic K^+^ levels caused by Kch activation).

We next examined how ionic control of porin permeability might influence susceptibility to antibiotics, many of which demonstrate porin-mediated entry into bacteria (*7*). Using single-cell fluorescence imaging, we observed reduced uptake of ciprofloxacin by OmpF, OmpC, and Kch deletion mutants (confirming porin-dependent uptake and the expected consequences of Kch regulation of porin activity), but no impact of deleting the efflux pump component TolC (*13*), indicating that, over the time course of these experiments, ciprofloxacin accumulation was dependent on influx rather than efflux (***Figure 4A***).

**Figure 4.**
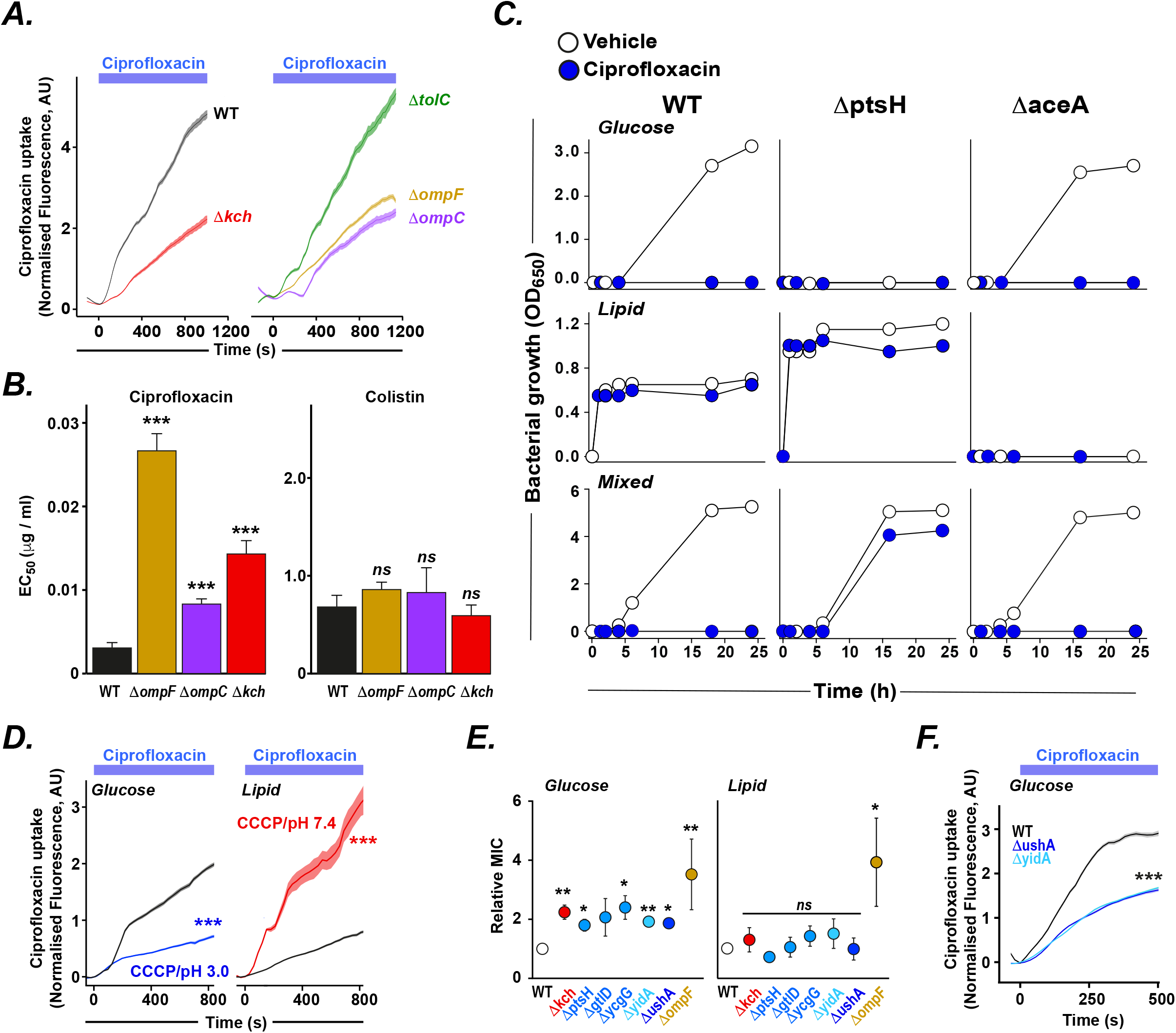
Metabolic control of porin permeability influences antibiotic resistance. (**A**) Accumulation of ciprofloxacin in individual bacteria over time (monitored by fluorescence imaging of type *E. coli* using a microfluidics platform) assessed for wild-type bacteria (WT; *black*), and isogenic knockouts for *kch* (*red*), the efflux pump component *tolC* (*green*), *ompF* (*yellow*) or *ompC* (*purple*). Fluorescence normalized to 0 s value. The data shown here (mean ± SEM) is representative of at least three independent experiments per condition with at least 50 individual bacteria imaged each time. \*\*\**p* < 0.001 (Student’s t-test). (**B**) EC50 values for wild-type *E. coli* (WT; *black*) and isogenic knockouts of *kch* (*red*), *ompF* (*yellow*) and *ompC* (*purple*) exposed to ciprofloxacin (*left*) or colistin (*right*). Data shown (mean ± SEM) is representative of at least three independent experiments performed at least in triplicate per condition. *ns* not significant, \*\*\**p* < 0.001 (Student’s t-test). (**C**) Bacterial growth (monitored by OD650) of wild type *E. coli* (WT’ *left*) or isogenic knockouts of *ptsH* (*middle*) or *aceA* (*right*) in glucose media (0.014 g/L DPPC + 0.3 g of Bacto Casitone and 0.5 mL of Tyloxapol; *top row*), lipid media (M9 + 1 g/L glucose; *middle row*), mixed media (0.014 g/L DPPC + 0.3 g of Bacto Casitone and 0.5mL of Tyloxapol + 1 g/L glucose; *bottom row*) in the presence of ciprofloxacin (0.0125 mg/mL; *blue*) or vehicle alone (*white*). (**D**) Accumulation of ciprofloxacin in individual wild-type *E. coli* over time (monitored as in (A)) during exposure to media containing glucose (description here; *left*) or lipid (description here; *right*) or during exposure to pH 3.0 media with CCCP (250 mM; *blue*) or pH 7.4 media with CCCP (250 mM; *red*). (**E**) Minimal inhibitory concentrations (MIC) of ciprofloxacin for wild type (WT, *white*) or isogenic knockouts for *kch* (*red*); *ptsH, gtlD*, or *ycgG* (*blue*); yidA (*light blue*), ushA (*dark blue*), or *ompF* (*yellow*) grown in glucose (*left*) or lipid (*right*) media (MIC values normalized to WT for each media). (**F**) Accumulation of ciprofloxacin within individual *E. coli* (monitored as in (A)) grown in glucose media (M9 + 4 g/L glucose) comparing wild type (WT, *black*), *ushA* (*dark blue*) *or yidA* (*light blue*) isogenic knockout strains. Data shown (mean ± SEM) is representative of at least three independent experiments per condition with at least 150 individual bacteria imaged each time. \*\*\**p* < 0.001 (Student’s t-test).

We found that ompF, ompC, and Kch knockout mutants showed increased resistance to ciprofloxacin (but not to colistin, an antibiotic that is porin-independent (*3, 56, 57*)), suggesting functional consequences of altered porin permeability (***Figure 4B***). We, therefore, wondered whether the recognized effects of different carbon sources on antibiotic susceptibility (*55, 58*– *62*) could be mediated through changes in porin permeability. We first confirmed that ciprofloxacin susceptibility was greater for *E. coli* grown in glucose compared to lipid media (***Figure 4C***). In order to understand the potential effects of different carbon sources on antibiotic stability, solubility, or availability, we examined the behaviour of mutants unable to grow in glucose (*ΔptsH*) or in lipid (*ΔaceA*) media. When grown in mixed media, wild-type bacteria and *ΔaceA* mutants demonstrated susceptibility to ciprofloxacin while *ΔptsH* mutants were found to be resistant (***Figure 4C***), implying that utilization of glucose leads to antibiotic susceptibility.

We observed that ciprofloxacin uptake was greater in bacteria grown in glucose than in lipid media but could be decreased in glucose media by increasing periplasmic H^+^ (by exposure to CCCP and pH 3 external media) and could be increased in lipid media by decreasing periplasmic H^+^ (by exposure to CCCP and pH 7.4 external media) (***Figure 4D***); findings that indicate that ionic regulation of porin function may mediate metabolic control of antibiotic susceptibility.

Since mutations in genes involved in central metabolism have been shown to cause antibiotic resistance in clinical and experimental settings (*59*), we wondered whether their effect might be mediated, at least in part, through changes in porin permeability. Deletion mutants for several genes previously implicated in drug resistance (*59*) showed increased ciprofloxacin resistance in glucose but not in lipid media (***Figure 4E***) and demonstrated reduced ciprofloxacin uptake when grown in glucose (***Figure 4F***), suggesting that these mutations prevent porin opening triggered by high metabolic activity.

Our findings, therefore, demonstrate that the permeability of porins in *E. coli*, and potentially other Gram-negative bacteria, can be dynamically regulated through changes in periplasmic H^+^ and K^+^, which in turn are influenced by the metabolic state of the cell via the action of the electron transport chain and the Kch potassium channel. Ionic control of porin permeability thus provides a mechanism by which nutrient uptake is linked to substrate utilization while preserving the periplasmic proton motive force. Our results also explain the recognized reduction in antibiotic susceptibility when bacteria are grown in lipid media and the impact of mutations in central metabolism genes on antibiotic resistance.

Our results suggest that therapeutic activation of Kch may enhance antibiotic accumulation within bacteria and could improve killing under conditions where lipids are utilized as the main carbon source, such as during intracellular infection (*63, 64*).

## Acknowledgements

We thank Jehangir Cama and the Keyser lab for help with the mother machine master template. We also help Emmanuel Derivery for help with the microfluidics facility. Funding: Supported by Wellcome Trust grants Investigator award 107032AIA (R.A.F., S.E.C.M, A.H., I.E.E.), the UK Cystic Fibrosis Trust [Innovation Hub grant 001 (S.E.C.M, A.H., T.L.B., R.A.F.); the NIHR Cambridge Biomedical Research Centre (R.A.F.); Botnar Foundation grant 6063 (S.E.C.M., R.A.F., T.L.B). Author contributions: R.A.F. conceived the project and wrote the manuscript; R.A.F. and S.E.C.M. designed the experiments. S.E.C.M., A.H., I.E.E, and D. S. performed the experiments; A.A. and T.L.B performed the molecular dynamics simulations. R. A. F. provided supervisory support. Competing interests: None.

## Supplementary Materials for

### MATERIALS AND METHODS

#### Bacterial strains and growth conditions

##### Bacterial strains

The following strains were from the KEIO collection (Horizon Discovery): *Escherichia coli* K12 BW25113 WT, *ΔompF, ΔompC, ΔompG, ΔnmpC, ΔphoE, Δkch, ΔclcB, ΔptsH, ΔaceA, ΔgtlD, ΔycgG, ΔyidA, ΔushA, ΔtolC*. The long-term stock was stored in 20% glycerol at -80 °C. Bacterial samples were stroked monthly in LB + agar with or without antibiotics in which kanamycin was used for selection at 50 µg/mL. Single colonies were resuspended in a 50 mL falcon tube with 10 mL fresh media as indicated. These cultures were incubated in glass test tubes with a breathable lid in a shaking incubator at 37°C in an orbital shaker at 300 rpm.

##### Media composition

The Super-Optimal Broth (SOB) or minimal medium 9 (M9) were used for each experiment. The super-optimal broth was prepared following a premix recipe (Fromedium, SOB01CFG). To prepare the minimal medium, we adapted the recipe described by Kotte(*65*): The medium contained the following components: Base salt solution (211 mM Na_2_HPO_4_, 110 mM KH_2_PO_4_, 42.8 mM NaCl, 56.7 mM (NH_4_)_2_SO_4_, autoclaved and prepared by the scientific facilities at LMB), 10 mL of trace elements (final concentration: 0.63 mM ZnSO_4_, 0.7 mM CuCl_2_, 0.71 mM MnSO4, 0.76 mM CoCl_2_, autoclaved), 0.1 mL 1 M CaCl_2_ solution (autoclaved, prepared by the scientific facilities at LMB) for a final concentration of 0.1 mM, 1 mL 1 M MgSO_4_ solution (autoclaved, prepared by the scientific facilities at LMB) for a final concentration of 1 mM, 2 mL of 500x thiamine solution (1.4 mM in ultrapure water from Milli-Q, Millipore and filter sterilized) and 0.6 mL 0.1 M FeCl_3_ solution (filter sterilized). The final volume was adjusted to 1 L with ultrapure water (Milli-Q, Millipore) lab water system (Milli-Q Advantage-10, Millipore). For each batch, the medium was filtered through 0.22 µm Millipore Stericup and split into 500 mL bottles. Before each experiment, the medium was supplemented with 1 g/L casamino-acids, 1 mM tryptophan and 0.5 g/L glucose unless the contrary was specified. Tryptophan stock was prepared at 50 mM in water from powder (Sigma Aldrich) and kept at 4 °C. Casamino acids (VWR Life Science) stock was prepared at 100 g/L in MiliQ water and kept at 4 °C.

##### Carbon sources

The carbon sources used in this work (glucose, fructose, acetate, fumarate, pyruvate) were acquired from Sigma-Aldrich in powder form. Then, stock solutions were prepared at a 100 g/L concentration in Milli-Q water. After adjusting the pH to 7.0, 100 mL aliquots were filtrated with a syringe filter, 0.22 µm. For the lipid media preparation, we used 14 mg/L 1,2-dipalmitoylphosphatidylcholine (DPPC), combined with 0.3 g/L casitone and 0.05% Tyloxapol to allow proper dilution.

#### Permeability quantification using flow cytometry

##### Tracer uptake

The method for permeability estimation in bacteria was derived from Jarzembowski, Hamilton and Jeon (*27, 33, 66*). Briefly, bacteria cultures were grown overnight at 37 °C in minimal medium M9, supplemented with casamino acids 1 g/L, tryptophan 1 mM and glucose 0.5 g/L. The next day, the source cultures were diluted to an OD∼0.05 in a 25 mL fresh medium and placed in a 250 mL conical flask. Glucose was not added to the medium for 2NBDG experiments. When the turbidity reached an OD of 0.1-0.25, the bacteria were split in a 96-well U-shape plate with 180 μL per well. Here, 10 μL of the treatment solution was added, and the plate was sealed with gas-permeable film (4titude, PCR0548) and returned to the incubator. After 30 min, 10 μL of the fluorescent tracer from the reservoir was added. Then, the concentration of the fluorescent tracer in the reservoir was diluted so we could keep the final volume constant.

After adding the fluorescent probe, the plate was returned to incubation for the specified time. Next, 200 μL were transferred to a V-bottom 96-well plate (Costar), and it was centrifuged 5 min x3900 rpm (Eppendorf 5810R, rotor S-4-104) and washed twice with PBS. Finally, samples were fixed and resuspended in PBS + 4% formaldehyde, stored at 4 °C and analyzed after 16 h. Then, we used the iCyt Eclipse (Sony) flow cytometer to read the fluorescence from all the samples. This device is equipped with 488 nm, 561 nm and 642 nm laser sources along with Hoechst and FITC filter sets.

For the data analysis, background fluorescence was subtracted and cell size effect was corrected by dividing tracer fluorescence by the forward scatter signal (*67*). For each experiment, the signal was normalized to WT strain or vehicle treatment.

##### Tracer uptake in different carbon sources

For the permeability estimation in different carbon sources, bacteria were grown as described before. When the cultures’ turbidity reached an OD of 0.1-0.25, the cultures were spun down 5 min x3900 rpm (Eppendorf 5810R, rotor S-4-104), washed with M9 and resuspended in 15 mL fresh medium. The different medium compositions were M9 + casamino acids 1 g/L, tryptophan 1 mM and glucose 0.5 g/L, M9 + 4 g/L (high) or 0.04 g/L (low) glucose, or M9 only (no carbon source). Next, the cultures were transferred to a 37°C water bath, and 400 μl samples were taken at the indicated timepoints. A 400 μl sample was transferred to an Eppendorf tube, and 4 μl Hoechst (1 mg/mL) was added to a final concentration of 1 μg/mL. The tubes were inverted 3 times and incubated at 37°C for 10 min. The samples were centrifuged for 30-sec max speed at 4 °C (Eppendorf centrifuge 5415 R), washed once and resuspended in 400 μl ice-cold PBS. Data analysis was performed as described in the previous section.

#### Microfluidics imaging

##### Mother machine design and assembly

We used the Mother Machine design described by Wang (*69, 70*). The microfluidic device was fabricated from an epoxy master template courtesy of Dr Jehangir Cama (University of Cambridge/University of Exeter, UK). The Mother Machine consists of a feed trench (50 µm x 100 µm x 30 mm) with many channels (1.4 µm x 1.4 µm x 25 µm) attached perpendicular to the trench.

For each batch of 12 chips, 50 mL of PMDS were mixed with a 5 mL curing agent (10:1) with vigorous stirring. The bubbles formed during the mixing were removed by vacuum degassing for 20 min until all air bubbles disappeared. Then, the gel was poured onto the master template and baked at 100 °C for 1 h. Subsequently, the chip was cut out around the wafer and prepared for bonding with the cover slide. Holes for inlets and outlets were punched using a sharpened 0.6 mm biopsy puncher (from Fisher Scientific), and the chip was cleaned with Scotch tape and 2-propanol. After drying all the excess isopropanol, the coverslip and PDMS with the features upwards were exposed to air plasma in a vacuum for 20 s at 0.4 mBar Oxygen (PlasmaPrep2, Gala Instrumente). This process activates the PDMS, which was put on a glass coverslip (24 mm x 50 mm, thickness 0.17 mm+/-0.005, Carl Roth). Finally, the chips were incubated overnight at 65 °C.

Once the chip was ready, we flushed the chamber with 2-iso-propanol and sonicated the chip for 30 min to remove all the debris (FB50, Fisher-Scientific). Then, we coated the internal chip surface to facilitate bacterial attachment with a passivation buffer (containing herring sperm DNA and BSA). The passivation buffer contained 10 mg/mL of Bovine Serum Albumin and 10 mg/mL of Salmond Sperm in a 3:1 ratio (*70*). The process was carried out overnight. Finally, the chips were stored in a dry cupboard for a week until required.

##### Trapping cells in the Mother machine

The bacteria preparation started with an overnight culture in the specified medium. Before injecting the cells into the microchip, 2 mL of cells were washed by centrifugation (Eppendorf 5810R) and resuspended in 2 mL of fresh medium. These cells were incubated for 10 min after and then centrifuged and concentrated to 100 µL. This high-density culture was injected into the microfluidic chip and set in a plate shaker at 37 °C for 10 min at 300 rpm (Grant-bio, PHMP). Once the cells were trapped in the channels, the chip was connected with an inlet and outlet tubing (Tygon ND-100-80 Medical Tubing - 0.5 mm ID, 1.52 mm OD). Next, we flushed the chip with plain M9 for 10 min so that any cells remaining in the main channel were removed.

##### 2NBDG uptake

2NBDG permeability was estimated by measuring the increase of fluorescent signal of *E. coli* cells trapped in the Mother Machine side channels. The recording set-up of the 2NBDG movies was done in a Nikon-N STORM, with a CCD camera (Andor-DU-897) and a 100x/1.49NA lens. The microscope had a closed environment chamber to maintain the temperature constant at 37 °C. For illumination, we used a 488-nm and 561-nm laser light (Aligent Technologies, MLC-400B). The 2NBDG fluorescent signal was captured with a dichroic mirror 525/50 (TRF49909), and for the 561-nm laser excitation experiments, we used a quad-band filter (97335) in combination with a 525/50nm filter.

##### Ciprofloxacin uptake

Cells were grown overnight in M9 + 1 g/L casamino acids + 0.5 g/L glucose + 1 mM tryptophan. The next day, cells were trapped in the mother machine as described in the previous section. Then, cells were resuscitated for 1-2h until growth started. Then, 12.5 µg/mL of ciprofloxacin was added to the medium and started flowing into the chamber. Ciprofloxacin uptake was measured in a Zeiss 780 microscope, equipped with a UV light source (DPSS 355 nm 60 mW, Coherent), which allowed ciprofloxacin imaging. This instrument was also equipped with a thermal isolation box to keep the temperature constant at 37 °C. The imaging was done with a 63x/1.4NA Oil lens. Ciprofloxacin fluorescence detection was done with a modified DAPI filter setting (435/65 nm)(*71*).

##### Ion sensor calibration

Cell trapping was done as described above. Cells were then resuscitated with M9 + 0.5 g/L glucose + 1 g/L casamino acids + 1 mM tryptophan for 90 minutes. Then, for pH calibrations, we switched the medium to M9 + 1 g/L glucose for 30 min and finally to PBS calibrated to a determinate pH with 1 g/L glucose. The pH selected points were 8.5 and 5.5. The same procedure was done by adding 250 µM CCCP to the medium to permeabilize the cells. In the case of the potassium calibration, we used a medium HEPES pH 7.4 with complementary concentrations of KCl and NaCl to keep the osmolarity constant at 150 mM. To permeabilize cells to the extracellular potassium, we treated cells with 100 µM valinomycin. The flowing conditions were kept at 0.15 mL/h, and during the switch, we increased the speed to 1 mL/h for 15 min. The illumination conditions for each sensor were the following: pelBCpHuji (Ex: 561 nm, Em: 620/60 nm), pHluorin (Ex: 488 nm, Em 525/50), ginko1 (Ex: 488 nm, Em 525/50), pelBCginko2 (Ex: 488 nm, Em 525/50), GCaMp6f (Ex: 488 nm, Em 525/50), QuasAr2 (Ex: 561 nm, Em 650/LP).

##### Ion sensor fluctuations

To observe the ion oscillations, we resuscitated cells with M9 + 0.5 g/L glucose + 1 g/L casamino acids + 1 mM tryptophan. The flow and temperature were kept constant at 0.150 mL/h and 37 °C. After 90 minutes, we started the recording process, which lasted until cells started dividing. For the data analysis, only 45 minutes before starting cell division were considered.

##### Carbon source switches

After trapping the cells into the chip, cells were washed with a plain M9 medium. Once the main channel was clear, the flow was kept constant at 0.15 mL/h for 30 minutes. At this point, we changed the syringe to the required carbon source and increased the flow to 1 mL/h. After flowing the medium for 2 minutes, we started the recording.

#### Agarose pad experiments

Cells were grown overnight in SOB + Ampicillin 100 µg/mL supplemented with 0.002 g/L arabinose and 20 µM retinal if required. The next day, cells were washed with M9 supplemented with the specified carbon source and resuspended for 10 min. Then 2 µL were transferred to the agar pad. Before starting the recording, pads were left drying for 5 min.

Agar pads were prepared on the same day of the experiment and discarded afterwards. The gel was composed of a 10 mL M9 medium with 1.5 % (wt/vol) low-melting agarose and the specified carbon source. We then proceed to dissolve the agarose by heating. Once the mixture was entirely homogeneous, the liquid was split into 3 mL portions onto a 35-mm petri dish (Falcon, ref: 353001). These plates were left to dry for 30 min. Once the agar solidified, we used a 6-mm biopsy puncher (Uni-Core, Harris UK) to cut out a single-use disk. We transferred 1 µL of cells on top of these agar disks, and after the drop was absorbed, the pad was moved onto a CELLview(tm) Dish with Glass Bottom (627870, Greiner). Recording started immediately.

#### Membrane voltage spike frequency

The QuasAr2 spike count was modelled with the glmmTMB R packages. The model considers the concentration of the carbon sources (in log scale) and corrects for the non-spiking cells. The model parameters are listed below.

|  | Estimate | Std. Error | Z-value | p-val (P(x>\|z\|)) |
| --- | --- | --- | --- | --- |
| *CSFructose* | 0.685776268 | 0.112229999 | 6.110454181 | **9.93E-10\*\*\*** |
| *CSFumarate* | 0.025517031 | 0.116852834 | 0.218368954 | 0.827141653 |
| *CSGlucose* | 0.781766915 | 0.116089923 | 6.734149658 | **1.65E-11\*\*\*** |
| *CSPyruvate* | 0.887781304 | 0.108779839 | 8.161266949 | **3.32E-16\*\*\*** |
| *Log(conc.)* | 0.013849301 | 0.005819853 | 2.379664808 | **0.017328392\*** |

#### MIC and EC50 measurements

Minimal inhibitory concentrations (MICs) and Effective concentration 50 (EC50), defined here as the drug concentration that induces a 50% maximal inhibitory effect on cell growth(*72*), were determined for *E. coli* according to the Clinical and Laboratory Standards Institute (CLSI) method M07-A9 (*73*). Briefly, *E. coli* strains were grown to an optical density (A600 nm) of 0.2–0.3 in liquid culture, and 1 × 10^5^ bacteria were added to each well of 96-well plates containing serial dilutions of the antibiotic in triplicate wells per condition and incubated at 37°C until growth was seen in the control wells. Then, the turbidity absorbance at 600 nm was measured with a ClarioStart (BGM LABTECH). Finally, the data were fitted to a dose-response model using a 4-parameter logistic regression equation (*74*). For the experiments in different carbon sources, cultures were grown in M9 + 4 g/L glucose or M9 supplemented with lipids (see above for full details).

#### Growth curves in the presence of ciprofloxacin

Individual *E. coli* colonies (WT, *ΔptsH* and *ΔaceA*) were resuscitated in 10ml of M9 media supplemented with 1 g/L glucose, M9 + lipid media (see Bacterial growth section for detailed composition) or mixed medium into 50 mL Falcon tubes. These cultures were incubated overnight at 37°C shaking. Note that the cultures grown in lipid media for slightly longer due to their slower growth rate in this carbon source. Next, the cultures were grown to an optical density (OD_650_) between 0.2-0.3. Finally, these cultures were diluted at 1:1000 and McFarland unit readings were taken at 1, 2, 4, 8, 16, and 24 hours.

#### Plasmid design and construction

The plasmids presented in this work were constructed using Gibson assembly (*75*). The plasmids pBAD_QuasAr2 (QuasAr2, #64134), pBAD_Ginko1(ginko1, #113111), and pKL004 (GCaMp6f, #98920), were obtained from the Addgene database (*48, 76, 77*). The plasmids for the expression of the pH sensor pHluorin were developed in previous work (*78*). Finally, the plasmids pBAD_ArchT, pBAD_pelBCpHuji and pBAD_pelBCginko2 were developed specifically for this work.

The primers were designed with the Primer3 algorithm available through the Benchling platform (*79*). The DNA material was amplified by PCR with the PrimeStart HS polymerase kit (Takara), following the manufacturer protocol and adjusting the annealing temperature to the suggested by the Benchling cloning algorithm. The amplification cycles were repeated 30 times using a C1000 Touch Thermal Cycler (BioRand).

The resulting DNA fragments were ligated with the following Gibson assembly kit. The concentration of the DNA templates was estimated with UV absorption using a NanoDrop 3300 (ThermoFisher), and the NEBiocalculator helped us estimate the required volumes for the Gibson assembly. All assemblies were carried out for 1h at 50 °C using the same thermocycler as the PCR amplification. Bacterial transformation of the assembled sequence was carried out through a heat shock using *E. coli* DH5α as an intermediate strain. Isolated plasmids were stored at -20 °C in Qiagen Elution Buffer.

##### pBAD_pelBCpHuji assembly

The periplasmic pH sensor was based on the pBAD-TOPO vector. This backbone was amplified with the primers pBAD_fwd (GCCACCCGCAGTTCGAAAAATAA<u>GTTTAAACGGTCTCCAGCTT</u>) and pBAD_rev (GCGGTCGGCAGCAGGTACTTC-AT<u>GGGTATGTATATCTCCTTCTTAAAG).</u> To this backbone, we added the *pelB* leader sequence. The *pelB* export signal sequence has proven to be successful in delivering recombinant proteins to the periplasm of *E. coli*(*80, 81*). This signalling sequence consists of 22 amino acids placed at the start codon. The pelBC fragment was amplified with the primers: pelB_fwd (TGCTGGCCGCCCAGCCGGCCATGGGGGGTTCTCATCATCA) and pelB_fwd (AAGCTGGAGACCGTTTAAAC<u>TTACTTGTACAGCTCGTCCATG)</u>. Then, we inserted the pH sensor. In order to measure the periplasmic pH accurately, we looked for a sensor with a wide dynamic range because the periplasmic pH could be up to 2 units lower than the cytoplasm, which means a pH 5 (*82, 83*). Since pHluorin fluorescence collapses below pH 6, we opted for pHuji with a working range between 5 and 9(*84*). For the amplification of the pHuji fragment, we used the primers: pelBCpHuji_fwd (TGCTGGCCGCCCAGCCGGCC<u>ATGGGGGGTTCTC ATCA-TCA)</u> and pelBCpHuji_rev (TGATGATGAGAACCCCCCAT<u>GGCCGGCTGGGCGGCC-AGCA)</u>. During the assembly process, we used *E. coli* DH5α, and after validating the sequence, we transferred the construct to an *E. coli* K12 BW25113 WT strain.

##### pBAD_pelBCginko2 assembly

We use the pelBC-pHuji plasmid to extract the backbone with the periplasmic export signal pelB. Since GINKO1 is derived from GFP, which is not stable in the periplasm, we designed a new variant based on sYFP for improved periplasmic stability. Thus, we amplified the potassium binding region of GINKO1 with the primers KBP_sYFP_fwd (ACAAGCTGGAATACAACTTC<u>CCGGACGGCCTGTTCAACTT</u>) and KBP_sYFP_rev (ATG-

TACACGTTGTGCGAGTT<u>CTCCAGCTCCTCGGGGATTC)</u> and inserted it between the 171 and 172 amino acids residues of sYFP. The sYFP was amplified with the primers pelBC_sYFP_fwd (AGAAGGAGATATACATACCC<u>ATGAAGTACCTGCTGCCGACCG</u>), pelBC_sYFP_rev (AAGTTGAACAGGCCGTCCGG<u>GAAGTTGTATTCCAGCTTGT</u>), end_sYFP_fwd (GAATCCCCGAGGAGCTGGAG<u>AACTCGCACAACGTGTACAT</u>), and end_sYFP_rev (AAGTGGAGACCGTTTAAAC<u>TTATTTTTCGAACTGCGGGTGGC</u>). This strategy was designed following the GINKO1 architecture. The final construct was amplified inside *E. coli* DH5α, and after validating the sequence, we transferred the construct to the *E. coli* K12 BW25113 strain.

Other genetic material: the plasmids pBAD_GINKO1 (Addgene code: #113111), pBAD_QuasAr2 (Addgene code: #64134) and GCaM-mRuby (also called pJMK0004, addgene code: #98920) were obtained via Addgene. Finally, the pBAD_pHluorin plasmid was developed in previous work (*85*).

##### pBAD_ArchT assembly

We used the pLenti-CaMKIIa-eArch 3.0-EYFP plasmid for the ArchT amplification (*78, 86*). For the PCR amplification, the primers ArchT_fwd AGGAGATATACATACCCATG<u>ATGGACCCAATTGCACTGCAGGCGGGG</u> and ArchT_rev AGCCAAGCTGGAGACCGTTTTTACAC<u>GGCGGCCGCAGGCTCCGGGGCTTCCGTA</u> were used. The backbone was derived from the pBAD_TOPO plasmid via digestion with the NcoI and PmeI restriction enzymes. Then, both fragments were ligated with the Gibson Assembly Kit.

#### Molecular Dynamic Simulations

The trimeric OmpC (PBD: 2J1N, resolution 2.0 Å, with 346 amino acid residues) and OmpF (PBD: 2OMF, resolution 2.40 Å, with 340 amino acid residues) were used in these simulation experiment. All simulations were performed by GROMACS v4.6 (www.gromacs.org) with CHARMM36 force fields for 100ns. Both systems were prepared using the CHARMM-GUI web interface. The OmpC 2J1N periplasmic side chains (E2, E43, E189, K6, K308, D7, D48, D135, D141, D268) were protonated based on pk_a_ value calculated using PROPKA3(*87*), whereas all other residues were set as a common state at physiological pH. The OPM server was used for orientation and positioning the protein in the membrane, and each system was embedded in a pre-equilibrated neutral zwitterionic lipid phosphatidylcholine (POPC) bilayer. The simulations were performed at constant pressure (1 atm) and temperature (300 K). The results were analyzed with GROMACS; the porin diameter was measured using HOLE (*88*), and images were prepared using PyMol.

---

**Figure S1.**
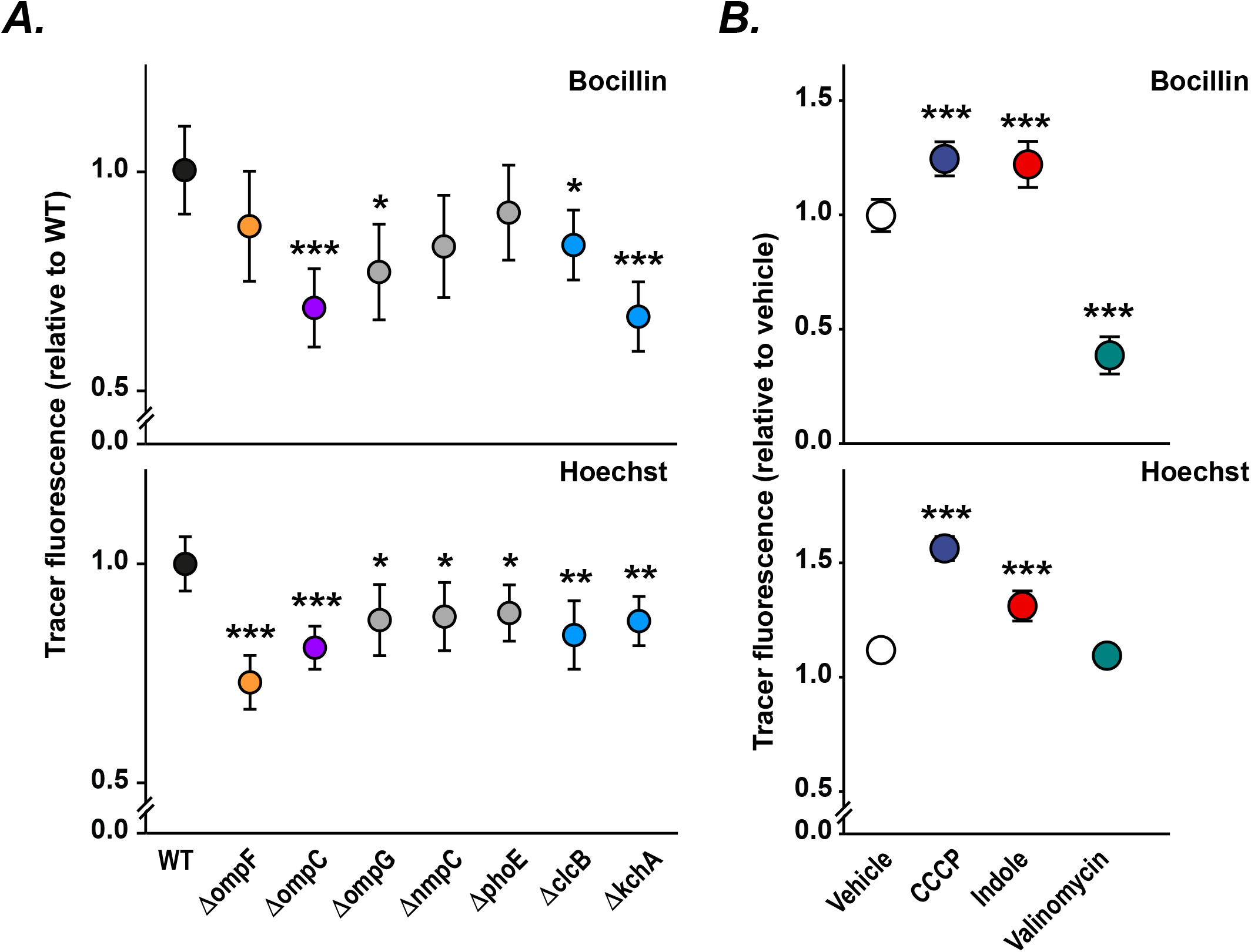
Accumulation of porin-dependent fluorescent tracers within *E. coli*. (**A**) Accumulation of bocillin (0.5 µg/mL; *top*) or Hoechst (1 µg/mL: *bottom*) in *E. coli* (quantified by flow cytometry) following 10 min incubation using wild type bacteria (WT; *black*) or isogenic knockouts for *ompF* (*yellow*) and *ompC* (*purple*), minor porins (*ompG, nmpC, phoE*; *grey*). (**B**) Accumulation of bocillin (0.5 µg/mL; *top*) or Hoechst (1 µg/mL: *bottom*) in wild type *E. coli* (quantified by flow cytometry) following treatment with the protonophores CCCP (concentration; *blue*) or Indole (concentration; *red*) or the potassium ionophore valinomycin (concentration; *green*), normalised to vehicle alone (*white*). Data (mean ± SEM) are representative of at least three independent experiments performed in at least triplicate. *p< 0.05, ** p< 0.01, *** p< 0.001 (Student’s t-test).

**Movie S1**. Representative single-cell fluorescence imaging of *E. coli* grown in a microfluidic perfusion system measuring 2NBDG accumulation over time in wild-type bacteria expressing empty vector (WT) or expressing the light-activated proton pump ArchT in the inner membrane in the presence of 541 nm light exposure.

**Movie S2**. Representative single-cell fluorescence imaging of *E. coli* WT and *Δkch* cells expressing the membrane potential reporter QuasAr2. Membrane depolarisation causes increased fluorescence.

